# Asthma Exacerbations: Integrative Analysis of miRNA Activity Using Single-Cell Transcriptomics

**DOI:** 10.64898/2026.07.31.741637

**Authors:** Parham Hadikhani, Xiting Yan, Geoffrey Chupp, Ga Young Ban, Shraddha Piparia, Michael McGeachie, Rinku Sharma, Scott T. Weiss, Louise C. Laurent, Alvin T Kho, Kelan G. Tantisira

## Abstract

**Background:** Asthma exacerbations are caused by dysregulated cellular interactions between airway and immune cell populations. Circulating microRNAs (miRNAs) are potential biomarkers for asthma exacerbations; however, their target airway cells remain poorly defined.

**Objective:** To identify the cell types that are regulated by the circulating microRNAs linked to asthma exacerbations and the extent to which the cells are regulated by miRNAs.

**Methods:** We integrated a curated panel of exacerbation-associated circulating miRNAs with single-cell RNA sequencing (scRNA-seq) profiles from induced sputum of 16 asthma patients and 8 healthy controls. Experimentally validated miRNA-target interactions were combined with cell-type-specific differential expression. Elastic Net regression and SHAP analysis quantified gene-level regulatory contributions, yielding a composite Regulation Strength metric. Findings were validated against four independent GEO datasets.

**Results:** Immune cells, including monocytes, dendritic cells, and macrophages, demonstrated the strongest statistically significant miRNA regulatory signals, in contrast to airway epithelial cells.hsa-miR-222-3p showed opposing regulatory effects in mature versus alveolar macrophages, indicating differentiation-state-dependent activity, while B_Plasma cells showed no detectable regulatory effect from any miRNA tested. Independent GEO validation confirmed higher expression of protective miRNAs (hsa-miR-126-3p, hsa-miR-146b-5p) in healthy individuals, consistent with prior CAMP cohort associations.

**Conclusion:** Circulating miRNAs show cell-type-specific regulatory activity, strongest in monocytes, dendritic cells, and macrophages. hsa-miR-222-3p showed opposing regulatory directions between macrophage subtypes, while B_Plasma cells showed no effect, validated across independent GEO cohorts.

## 1 Introduction

Asthma is a chronic inflammatory airway disease characterized by variable airflow obstruction, airway hyperresponsiveness, and recurrent exacerbations [1, 2, 3, 4]. Exacerbations represent acute periods of worsening respiratory symptoms that can be life-threatening and often arise in response to allergens, viral infections, or environmental triggers [5, 6]. Although these are clinically well recognized, the specific cellular mechanisms that drive exacerbation-associated inflammation remain insufficiently understood, and this gap limits the development of targeted therapeutic strategies [7].

miRNAs are small non-coding RNAs that regulate gene expression at the post-transcriptional level and influence a wide range of immune and epithelial cell functions [8]. Circulating miRNAs detectable in serum or plasma have emerged as promising biomarkers of asthma severity and exacerbation risk. miRNA mediate long range cell-cell communication. However, an important unanswered question is which lung cell populations are directly targeted by these circulating miRNAs during exacerbations and how miRNA-mediated regulation contributes to airway inflammation.

scRNA-seq provides high-resolution gene expression profiles from thousands of individual cells and has transformed our understanding of airway biology by revealing previously unrecognized cellular heterogeneity and dynamic disease-associated cell states [9, 10]. These data offer a unique opportunity to link systemic molecular signals to specific airway cell types. However, mature miRNAs lack poly(A) tails [11] and are extremely short (22 nucleotides) [12], which prevents efficient capture and amplification by standard scRNA-seq protocols. This limitation prevents direct measurement of miRNA expression in single cells and necessitates computational strategies to infer miRNA activity from mRNA-based single-cell transcriptomes.

To address this challenge, we developed a computational approach that analyzes a curated panel of circulating miRNAs associated with asthma exacerbations, including hsa-miR-206, hsa-miR-223-5p, hsa-miR-146b-5p, hsa-miR-342-3p, hsa-miR-222-3p, hsa-miR-30e-3p, hsa-miR-126-3p, and hsa-miR-126-5p [13]. By integrating these circulating miRNAs with scRNA-seq profiles from induced sputum samples of asthma patients and healthy controls [14], we systematically identified the airway cell types predicted to be influenced by each miRNA. Using experimentally validated miRNA-target interactions together with machine learning-based modeling, we infer miRNA regulatory activity at cell-type resolution and characterize the molecular effects that may contribute to exacerbation-associated inflammation.

This integrative approach provides a mechanistic link between circulating miRNAs and specific lung cell populations. Our study reveals cell-type specific miRNA regulatory patterns in asthma and identifies the primary cellular targets of exacerbation-associated miRNAs.

## 2 Methods

### 2.1 Overview

We developed a computational approach to identify the cellular targets and regulatory effects of circulating miRNAs during asthma exacerbations, integrating experimentally validated miRNA-target interactions, single-cell transcriptomic profiles, and machine learning-based modeling to infer miRNA regulatory activity at cell-type resolution (Fig. 1). The analysis proceeded in three stages: (i) integration of circulating miRNAs with cell-type-specific gene expression, (ii) inference of miRNA regulatory activity using a regression-based approach, and (iii) interpretation and validation of the inferred regulatory relationships. Full parameter settings, software versions, and formulae are provided in Supplementary Methods.

**Figure 1.**
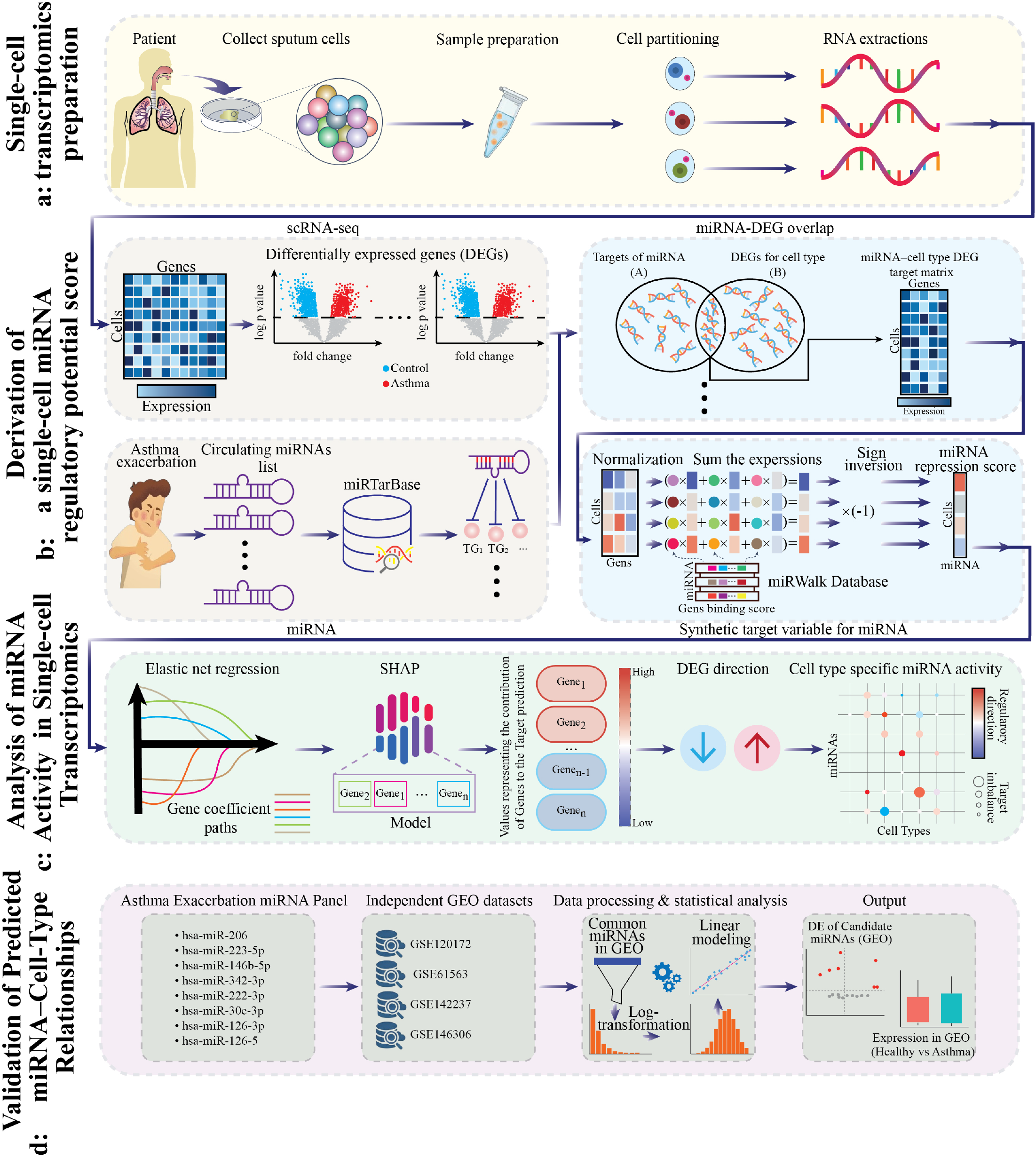
Overview of the computational approach integrating circulating miRNAs with scRNA-seq data. The workflow includes single-cell data preparation, derivation of cell-type–specific miRNA regulatory potential scores, and analysis of miRNA activity using Elastic Net modeling and SHAP interpretability, followed by validation of predicted miRNA–cell-type relationships using experimentally supported miRNA–target interactions and cell-type–specific gene expression profiles.

### 2.2 Data Sources

A panel of eight circulating miRNAs previously associated with asthma exacerbations (hsa-miR-206, hsa-miR-223-5p, hsa-miR-146b-5p, hsa-miR-342-3p, hsa-miR-222-3p, hsa-miR-30e-3p, hsa-miR-126-3p, and hsa-miR-126-5p) was curated from a serum-based study of the Childhood Asthma Management Program (CAMP) cohort [13], in which logistic regression identified miRNAs predictive of future exacerbations. Experimentally validated miRNA-gene interactions were retrieved using the multiMiR R package [15] (miRecords, TarBase, miRTarBase); a gene was retained as a target only if supported by at least two independent databases or experimental methods. scRNA-seq data were obtained from induced sputum samples of 16 asthma patients and 8 healthy controls [14] (37,565 cells; median 1,203 genes and mean 4,186 UMIs per cell). Clustering was performed using Seurat [16], yielding 28 clusters annotated into 19 cell populations based on canonical markers and correlation with the Human Primary Cell Atlas [17]. Cell-type-specific differential expression between asthma and control samples was computed using the FindMarkers function in Seurat (min.pct = 0.25, log_2_FC threshold = 0.25; significance defined as *p* < 0.05 and log_2_FC > 0.5).

### 2.3 miRNA Repression Score and Regulatory Modeling

For each miRNA, validated target genes were intersected with cell-type-specific differentially expressed genes (DEGs). Binding probabilities for miRNA-gene pairs, obtained from the miRWalk 3.0 database [18], were used as gene-level weights to compute a per-cell miRNA repression score, serving as a proxy for miRNA activity and as the response variable in downstream modeling (see Supplementary Methods for the full formula). For each miRNA-cell-type pair, Elastic Net regression [19] was trained using standardized target-gene expression as predictors and the repression score as the outcome, with five-fold cross-validation used to select regularization parameters. SHapley Additive exPlanations (SHAP) [20] were then applied to quantify each gene’s contribution to the predicted repression score, and combined with differential expression direction to infer whether a miRNA was preferentially repressing asthma-upregulated or asthma-downregulated genes.

### 2.4 Regulation Strength Metric

To quantify the magnitude and direction of miRNA regulation within each cell type, we defined a composite Regulation Strength metric integrating SHAP-derived gene contributions with differential expression direction (formula in Supplementary Methods). A positive value indicates preferential repression of asthma-downregulated genes (increased miRNA activity), while a negative value indicates preferential repression of asthma-upregulated genes (reduced miRNA activity); the magnitude reflects the strength of this directional bias. Statistical robustness was assessed using 95% bootstrap confidence intervals from 1,000 cell-resampling iterations (Supplementary Table 1).

### 2.5 External Validation

To validate the inferred regulatory relationships, four independent GEO datasets (GSE120172 [21], GSE61563 [22], GSE142237 [23, 24], and GSE146306 [25]) were analyzed using the GEOquery package [26] and the limma framework [27], with log_2_ fold changes and Benjamini-Hochberg-adjusted *p*-values [28] computed for each candidate miRNA. Consistent differential expression patterns across datasets were considered supportive of the predicted miRNA-cell-type regulatory relationships.

## 3 Results

### 3.1 Cell-Type–Specific Patterns of miRNA Activity

In Fig. 2, cell-type-specific regulatory effects mediated by miRNAs during asthma exacerbation are depicted based on regulation score, spatial position of the cell type, and statistical significance through miRNA targeting activity inferred from single-cell mRNA expression profiles.

**Figure 2.**
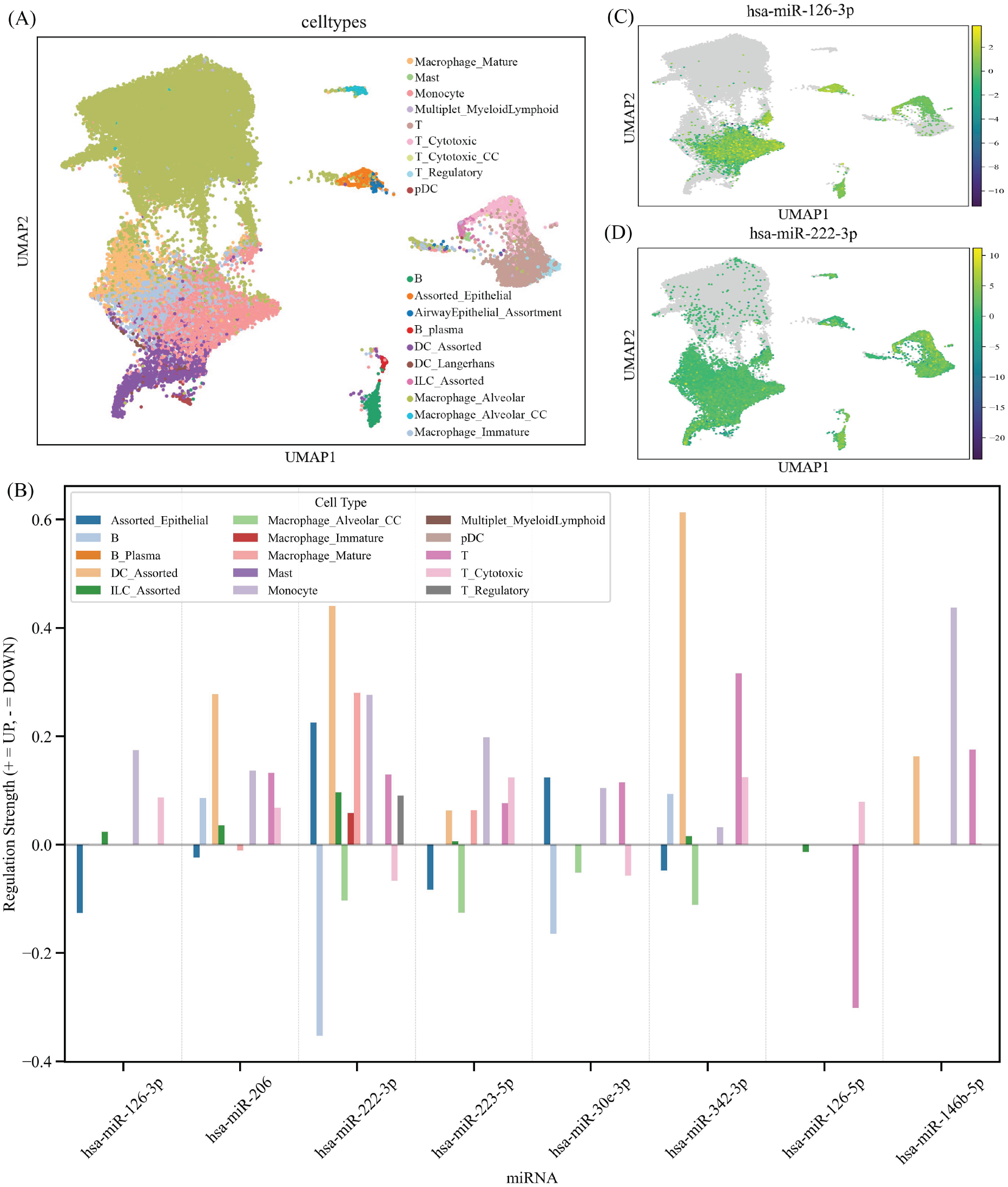
miRNA regulation in cell type-specific manner in asthma exacerbation. (A) UMAP representation of the induced sputum cells divided into 19 distinct cell populations. (B) Regulation Strength of eight miRNAs in 15 airway and immune cell populations. Negative numbers represent the bias towards the repression of upregulated genes, while positive numbers indicate a preference toward the repression of downregulated genes in asthma. Regulation strength is a measure combining SHAP importance with differential expression directionality. (C-D) Maps of miRNA activity significance within single-cell data for hsa-miR-126-3p (C) and hsa-miR-222-3p (D). The miRNA activity significance is defined as−log_10_(*p*-value), estimated by null model based on the permutation of the weight-target gene pairing. Higher values denote the increased likelihood of the non-random miRNA regulation. The cells colored in grey are omitted from the analysis due to lack of gene overlap. Here, the two miRNAs show distinct activity patterns, with hsa-miR-222-3p exhi7biting broader and more intense significance scores across myeloid, lymphoid, and epithelial compartments, while hsa-miR-126-3p activity is predominantly restricted to myeloid clusters.

By applying our approach, we analyzed induced sputum from asthmatic patients and healthy individuals based on single-cell RNA-seq, which included 37,565 total cells in all samples. After data quality control and preprocessing, the average number of unique molecular identifiers (UMIs) was 4,186, and the median number of genes detected per cell was 1,203.

Integrated clustering and uniform manifold approximation and projection (UMAP) analysis [29] resolved the dataset into 28 transcriptionally distinct clusters, which were subsequently annotated into 19 canonical cell populations based on established marker gene expression. Fig. 2(A) shows the UMAP representation of all cells, illustrating the separation of major airway and immune populations, including myeloid, lymphoid, and epithelial compartments. The cellular composition was predominantly of myeloid origin, with Macrophages representing the largest fraction (71.8%). This macrophage population included alveolar macrophages (accounting for approximately 81% of the total macrophage lineage) and several subclusters. Monocytes (12.7%) and dendritic cells (5.1%) were also identified, making up the remaining major myeloid populations. The lymphoid compartment, comprising T lymphocytes, innate lymphoid cells, and B lymphocytes, accounted for 9.3% of the total cells. Airway epithelial cells were also captured but represented a small fraction of the overall population (0.99%). To identify cell populations contributing meaningful miRNA-mediated regulatory signals, we evaluated the overlap between predicted miRNA target genes and cell-type–specific DEGs, as shown in Supplementary Fig. 1-4. Several annotated populations exhibited no detectable overlap and were therefore excluded, resulting in a refined set of 15 cell populations used for downstream integrative modeling.

Fig. 2(B) shows the direction and magnitude of regulatory influence for eight miRNAs across these 15 cell populations, quantified using the Regulation Strength metric described in Section 2.4. To assess the statistical robustness of these signals, 95% bootstrap confidence intervals were derived from 1,000 cell-resampling iterations; pairs with bootstrap *p* < 0.05 were considered statistically significant. Complete bootstrap statistics for all miRNA-cell type pairs are provided in Supplementary Table 1. Substantial cell-type variability in regulatory direction was observed. Notably, Assorted_Epithelial cells, a structurally central population in asthma pathobiology, exhibited Regulation Strength trends for multiple miRNAs, including hsa-miR-126-3p, hsa-miR-222-3p, hsa-miR-223-5p, and hsa-miR-342-3p, though none reached bootstrap significance (range:*p*= 0.054 to 0.658). Among these, hsa-miR-222-3p displayed the largest positive Regulation Strength, indicating preferential repression of transcripts already downregulated in asthma, while hsa-miR-126-3p, hsa-miR-223-5p, and hsa-miR-342-3p showed negative trends consistent with preferential repression of asthma-upregulated genes. hsa-miR-206 and hsa-miR-30e-3p showed near-zero non-significant signals in this compartment (*p* = 0.632 and *p* = 0.658, respectively).

Beyond epithelial cells, the strongest and most statistically robust regulatory signals were concentrated in innate immune populations. In dendritic cells (DC_Assorted), hsa-miR-222-3p (*p* = 0.001), hsa-miR-342-3p (*p* = 0.001), hsa-miR-206 (*p* = 0.001), hsa-miR-223-5p (*p* = 0.001), and hsa-miR-146b-5p (*p* = 0.004) all showed significant positive Regulation Strength. In Monocytes, hsa-miR-146b-5p, hsa-miR-126-3p, hsa-miR-206, hsa-miR-30e-3p, and hsa-miR-223-5p (all *p* = 0.001), as well as hsa-miR-342-3p (*p* = 0.004), were statistically significant, while hsa-miR-222-3p did not reach significance (*p* = 0.338). In Macrophage_Mature, hsa-miR-222-3p showed robust positive Regulation Strength (*p* = 0.001). Notably, in Macrophage_Alveolar_CC, hsa-miR-222-3p exhibited a significant negative Regulation Strength (*p* = 0.015), indicating a directionally opposite regulatory pattern compared to Macrophage_Mature and suggesting macrophage-subset-specific miRNA activity. Together, these myeloid populations were identified as the major cellular targets of statistically supported miRNA dysregulation in asthma.

T and cytotoxic T cells (T_Cytotoxic) also showed several significant regulatory signals. In T cells, hsa-miR-342-3p, hsa-miR-223-5p, hsa-miR-206, hsa-miR-30e-3p, and hsa-miR-126-5p were all significant (all *p* = 0.001). In T_Cytotoxic cells, hsa-miR-126-5p (*p* = 0.001), hsa-miR-206 (*p* = 0.001), hsa-miR-223-5p (*p* = 0.002), and hsa-miR-146b-5p (*p* = 0.003) were significant.

In contrast, B_Plasma cells showed no statistically meaningful regulatory signals across all tested miRNAs (all *p* = 1.000), indicating negligible miRNA regulatory activity in this compartment.

To validate and spatially contextualize these regulatory patterns, Figs. 2(C and D) show permutation-based significance scores for individual miRNAs projected onto the UMAP embedding, with Fig. 2(A) serving as the cell-type reference. For each cell, observed miRNA activity was compared against a null distribution generated by permuting the association between target genes and their SHAP-derived weights, the resulting p-values were transformed to −log_10_(*p*), with higher values denoting stronger statistical significance. The two miRNAs selected for spatial visualization, hsa-miR-126-3p (Fig. 2(C)) and hsa-miR-222-3p (Fig. 2(D)), illustrate markedly different activity profiles across the UMAP landscape, reinforcing the cell-type-specific directionality observed in Fig. 2(A). hsa-miR-126-3p activity was largely concentrated in the myeloid-dominant lower clusters, with moderate significance scores (−log_10_(*p*) up to approximately 2.5), and comparatively sparse signal in lymphoid and epithelial regions. By contrast, hsa-miR-222-3p exhibited substantially broader and more intense activity, with significance scores reaching −log_10_(*p*) ≈ 10 and spanning both myeloid and immune compartments. Notably, the elevated hsa-miR-222-3p signal extended into cell clusters corresponding to macrophage and monocyte populations, consistent with its large positive Regulation Strength in these lineages shown in Fig. 2(A). Together, these spatial maps demonstrate that the inferred regulatory relationships are statistically robust at the level of individual cells, with miRNA activity that is both cell-type-selective and non-random.

### 3.2 Association of Circulating microRNAs with Asthma Exacerbation Risk

We examined the clinical association between a previously reported panel of circulating miRNAs and the risk of asthma exacerbation. This panel was originally identified in the CAMP cohort, where circulating miRNAs were quantified directly in serum by TaqMan qPCR and evaluated by logistic regression for their relationship to future exacerbation risk [13]. In that study, eleven circulating miRNAs were significantly associated with exacerbation outcomes.

Eight of these miRNAs showed statistically significant inverse associations (odds ratio, OR < 1), suggesting that higher circulating levels are linked to a lower risk of exacerbation. The strongest associations were observed for hsa-miR-206 (OR: 0.60, 95% CI: 0.42–0.83, p = 0.004), hsa-miR-146b-5p (OR: 0.66, 95% CI: 0.48–0.89, p = 0.007), and hsa-miR-222-3p (OR: 0.70, 95% CI: 0.52–0.93, p = 0.02). Additional miRNAs with significant inverse associations included hsa-miR-223-5p, hsa-miR-126-5p, hsa-miR-30e 3p, hsa-miR-126-3p, and hsa-miR-342-3p. These previously reported findings guided the selection of circulating miRNAs for downstream cell-type–specific regulatory modeling in the present study.

### 3.3 Cell-Type–Specific Targeting of Circulating miRNAs

Fig. 3 shows the number of model-selected target genes for each miRNA-cell-type combination. Each value indicates how many genes the Elastic Net model predicted to be regulated by a given miRNA within that specific cell type. These counts reflect is based only on those genes which have been experimentally validated and which are differentially expressed in that particular cell population, resulting in smaller but more biologically relevant, cell-type-specific subsets. Rows correspond to the cell populations identified from scRNA-seq analysis, and columns correspond to the eight circulating miRNAs associated with asthma exacerbation risk.

**Figure 3.**
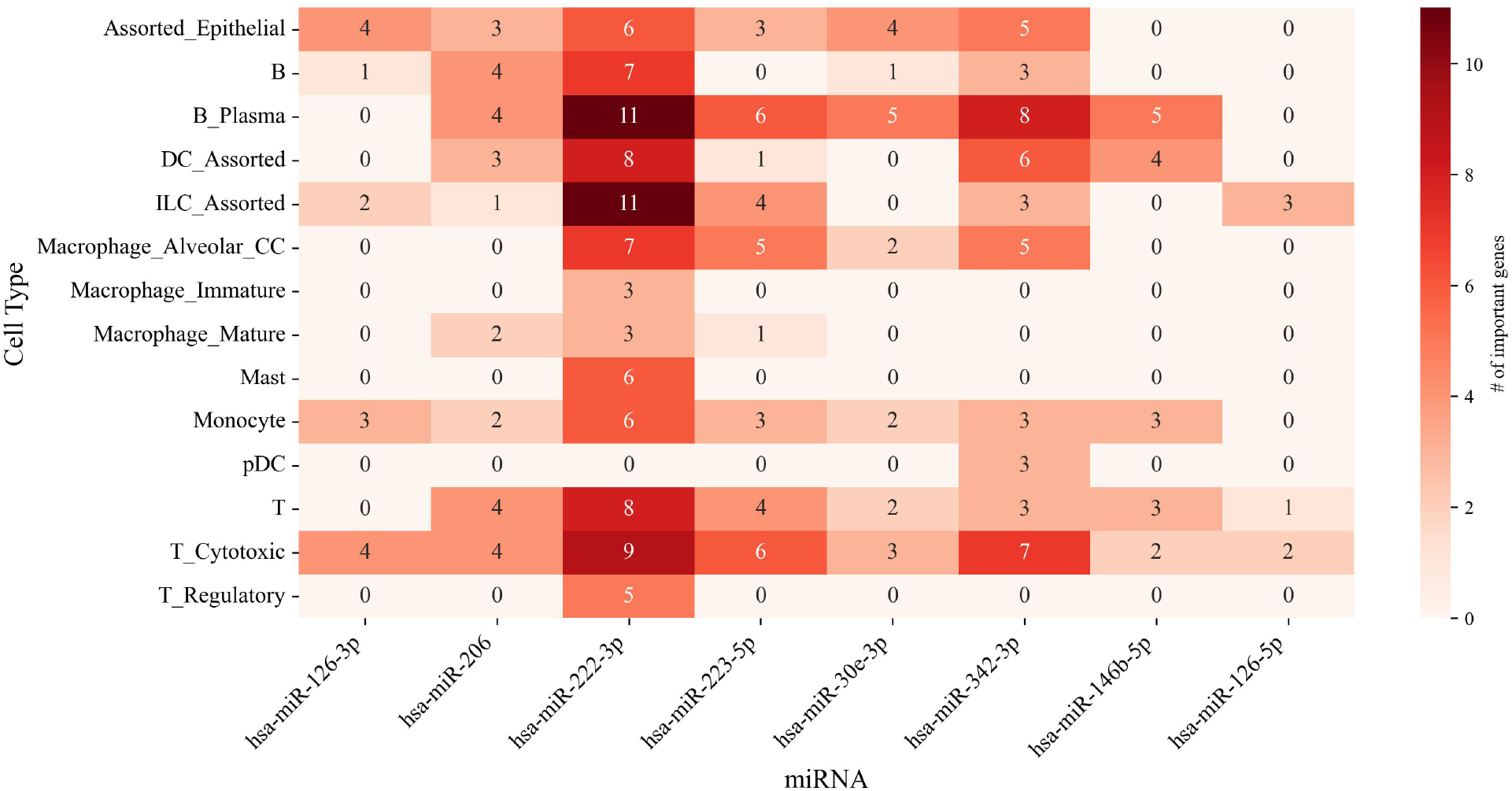
Number of model-selected target genes for each miRNA-cell-type pair identified by ElasticNet regression.

The resulting matrix reveals substantial variation in the number of selected targets across both miRNAs and cell types. Several miRNA–cell-type pairs show relatively large sets of informative genes. For example, hsa-miR-342-3p displays a high number of selected targets in B_Plasma cells, while hsa-miR-222-3p exhibits an enriched set of targets in innate lymphoid cells (ILC_Assorted). In contrast, certain cell types, including mature macrophage subsets, show consistently low numbers of selected targets across most miRNAs. Additionally, miR-126-5p had minimal differentially expressed target genes within the sputum cells.

### 3.4 Inferred Cell-Type–Specific miRNA Regulatory Activity

Fig. 4 visualizes the regulatory influence of circulating miRNAs across immune and epithelial cell types. Each dot encodes two pieces of information: dot size represents the number of target imbalance, calculated as the absolute difference between upregulated and downregulated target genes (|up_count − down_count| + 1), with larger dots indicating a stronger, directionally consistent effect of a miRNA on its targets within a given cell type. Dot color reflects the “target gene direction, defined as the difference between the weighted sums of down-versus up-regulated target genes. Red indicates that the miRNA acts as an activator (greater influence toward upregulated targets), purple indicates a suppressive effect (greater influence toward downregulated targets), and white indicates balanced or neutral regulation.

**Figure 4.**
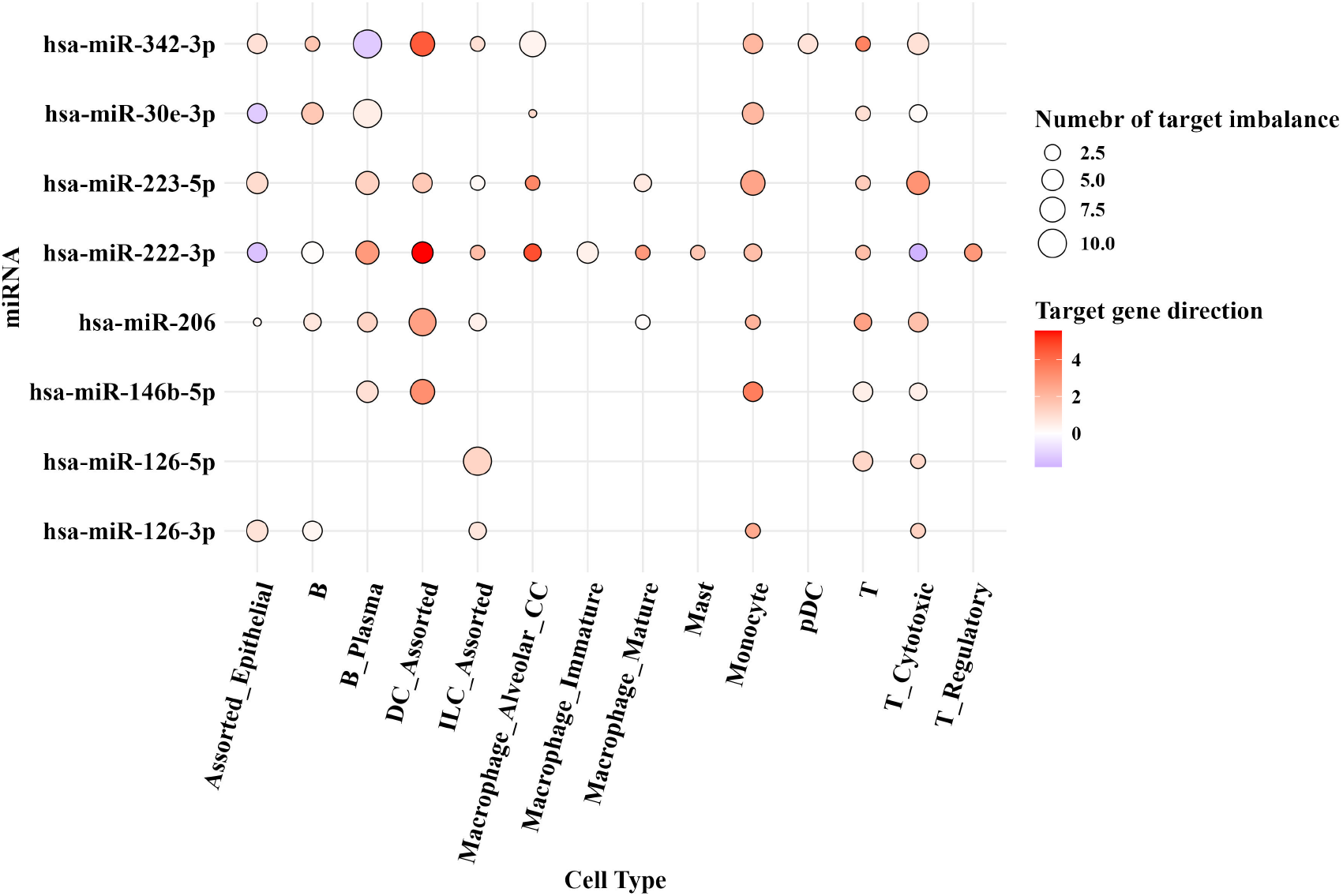
Point size reflects the magnitude of each miRNA's influence based on SHAP-derived target contributions, and point color indicates the dominant direction of target regulation in asthma. Cell type labels on the x-axis represent distinct populations identified by scRNA-seq clustering.

The targeting activity of individual miRNAs was highly cell type-specific. The most directionally heterogeneous miRNA is hsa-miR-222-3p, as it exhibits the largest dot value in B_Plasma with a moderate activating direction. In contrast, DC_Assorted shows a smaller but strongly activating dot (vibrant red), while Assorted_Epithelial indicates a suppressive effect of this miRNA on those structures. hsa-miR-342-3p also presents its largest activation values in B_Plasma and Macrophage_Alveolar_CC, with an overall mild activating direction. Notably, DC_Assorted exhibits a smaller dot but suggests strong activation with good directional consistency despite the low magnitude. hsa-miR-146b-5p shows its strongest activation signal in DC_Assorted (moderate), while Monocyte presents a similar but weaker signal, and B_Plasma shows moderate activation as well. hsa-miR-223-5p is primarily activating in Monocyte and T_Cytotoxic (moderate activation direction), with neutral activation observed in DC_Assorted and B_Plasma. hsa-miR-206 exhibits its strongest activation in DC_Assorted, along with mild activation signals in B_Plasma and T_Cytotoxic. hsa-miR-30e-3p exhibited a broadly diffused but consistent weak activation effect, with its maximum value in B_Plasma and weak activation signals in B cells, Monocyte, and Assorted_Epithelial. The hsa-miR-126-3p molecule had its highest value in Assorted_Epithelial and was exhibiting mild activation effects there, with weak activation found in B cells, ILC_Assorted, and T_Cytotoxic. The most restricted distribution was that of hsa-miR-126-5p, which only activated ILC_Assorted, T cells, and T_Cytotoxic.

### 3.5 Validation of Cell-Type-Specific miRNAs in Asthma

To validate the cell-type–specific miRNAs predicted from our integrated scRNA-seq and circulating miRNA analysis, we examined their expression in independent asthma datasets from GEO. We assessed whether these miRNAs were detectable, differentially expressed between asthma patients and healthy controls, and consistent with the inferred cell-type regulatory patterns.

We analyzed multiple GEO datasets encompassing asthma-relevant sample types, including bronchial epithelial cells, sputum, plasma, and peripheral immune cells (Table 1). For each dataset, miRNA expression values were extracted, samples were annotated with phenotype metadata, and differential expression was evaluated using the limma framework. Volcano plots highlighted significantly up or downregulated miRNAs (Supplementary Figs. 5–6) and Expression differences were visualized with boxplots (Supplementary Figs. 7–8).

**Table 1.** Summary of GEO Datasets Used for Validation.

| GEO Dataset | Disease Context | Sample Type / Cell Type | Pathway / Mechanism |
| --- | --- | --- | --- |
| GSE120172 [21] | Pediatric dust-mite asthma | Bronchial epithelial cells (HBE), plasma | PI3K-AKT |
| GSE61563 [22] | Chronic asthma | Bronchial epithelial cells (BEC) | Cytokine regulation |
| GSE142237 [23, 24] | Steroid-naïve asthma | Bronchial epithelial cells (BEC) | circRNA-miRNA-mRNA networks, IL-17 |
| GSE146306 [25] | Severe asthma | Sputum cells, various immune cells | miRNA-mRNA networks, TLR, Th17 |

<sup>a</sup>Pathway and mechanism information obtained from the original publications associated with each GEO dataset.

Differential expression analysis revealed consistent trends aligned with the predicted regulatory roles of the candidate miRNAs. For example, in GSE120172, hsa-miR-126-3p and hsa-miR-146b-5p were significantly higher in healthy controls (P = 0.0115 and 0.0296, respectively), consistent with their previously reported protective associations in the CAMP cohort [13], whereas hsa-miR-206 showed a modest increase in asthma (P = 0.0566). In GSE142237, hsa-miR-342-3p was significantly lower in asthma (P = 0.0295), supporting its predicted immunoregulatory function. Other miRNAs, including hsa-miR-30e-3p and hsa-miR-206, exhibited trends consistent with predictions but did not reach statistical significance, likely due to sample size limitations or cellular heterogeneity (Table 2).

**Table 2.** Differential Expression of Candidate miRNAs Across GEO Datasets.

| miRNA | Dataset | logFC | P-value | Trend (Asthma vs Healthy) |
| --- | --- | --- | --- | --- |
| hsa-miR-126-3p | GSE120172 | 2.266 | 0.0115 | Higher in Healthy (significant) |
| hsa-miR-146b-5p | GSE120172 | 1.732 | 0.0296 | Higher in Healthy (significant) |
| hsa-miR-206 | GSE120172 | -0.299 | 0.0566 | Higher in Asthma (marginal) |
| hsa-miR-342-3p | GSE61563 | 3.054 | 0.0589 | Higher in Asthma (marginal) |
| hsa-miR-342-3p | GSE142237 | -15.832 | 0.0295 | Lower in Asthma (significant) |
| hsa-miR-30e-3p | GSE146306 | -0.328 | 0.0530 | Higher in Asthma (marginal) |
| hsa-miR-206 | GSE146306 | -0.375 | 0.0757 | Higher in Asthma (marginal) |
| hsa-miR-146b-5p | GSE142237 | 4.372 | 0.0769 | Higher in Healthy (marginal) |
| hsa-miR-342-3p | GSE146306 | -0.567 | 0.0908 | Higher in Healthy (marginal) |

Cross-referencing these expression patterns with predicted cell-type specificity, as shown in Table 3, revealed general concordance with model predictions. For example, hsa-miR-126-3p, enriched in epithelial, DC, and select immune populations including ILC, B, monocyte, and T cytotoxic cells, was higher in healthy samples, whereas hsa-miR-206, broadly expressed across B cells, monocytes, epithelial, DC, ILC, macrophage, and T cells, was elevated in asthma samples.

**Table 3.** Predicted Cell-Type Specificity and Expression Trends.

| miRNA | Predicted Cell Types | Trend ($P < 0.05$) |
| --- | --- | --- |
| hsa-miR-126-3p | ILC, B, Monocyte, Epithelial, DC, T Cytotoxic | Higher in Healthy |
| hsa-miR-206 | B Plasma, Epithelial, B, Monocyte, T, DC, ILC, Macrophage | Higher in Asthma |
| hsa-miR-30e-3p | Epithelial, DC, B Plasma, Monocyte, T, Macrophage | Higher in Asthma |
| hsa-miR-342-3p | DC, Macrophage, Monocyte, T, pDC, B Plasma, Epithelial, T Cytotoxic | Lower in Asthma |
| hsa-miR-146b-5p | B Plasma, DC, Monocyte, Macrophage, T Cytotoxic, T | Higher in Healthy |

## 4 Discussion

In this study, we combined miRNA profiling with single-cell gene expression data in sputum samples to reveal cell type-specific miRNA regulation linked to exacerbation-associated miRNAs. We show that these miRNAs exhibit unique cell-type specific regulatory functions in epithelial and immune cells via SHAP scores, rather than being uniformly regulated, suggesting that each miRNA contributes uniquely to the mechanism of asthma exacerbations.

### 4.1 Cell-type-specific regulatory influence of circulating miRNAs

The results of our cell type-specific analysis demonstrated very heterogenous regulatory signatures, which suggest that each circulating miRNA possesses its own transcriptional signature. The role of circulating miRNAs is to regulate cells specifically rather than to serve as universal regulators within the body system.

The monocyte, macrophage, dendritic, and cytotoxic/ regulatory T-cell lineage were identified as major players affected by circulating miRNA. The cellular lineages showed high Regulation Strength for several miRNAs, indicating susceptibility to miRNA-mediated targeting relevant to asthma exacerbation pathobiology, as inferred from stable asthma patient samples. In the category of structural cells, the Assorted_Epithelial cell lineage showed high regulatory significance, signifying the importance of epithelial barrier communication in asthma [30].

Directional miRNA targeting activity patterns were yet another confirmation of this dependence of effect on context. For example, there was reduced targeting activity for hsa-miR-222-3p by some immune subsets, which resulted in up-regulation of its target genes, due to the lack of suppression by the miRNA in those cell populations. In turn, miRNAs such as hsa-miR-126-3p and hsa-miR-342-3p were found to be more selective and act primarily on the lineage-specific targets. Of the total pool of eight miRNAs, hsa-miR-126-5p turned out to have particularly limited targets and activity, being effective only in the case of ILC_Assorted, T, and T_Cytotoxic cell types. Thus, miRNAs may affect each individual cell type based on its microenvironment differently, amplifying or reducing asthmatic signals, accordingly.

### 4.2 Implications of Divergent miRNA Regulatory Signals in Cell Types During Asthma Exacerbation

The variability of Regulation Strength scores in epithelial and immune cell types emphasizes the heterogeneous impact of circulating miRNAs on asthma exacerbations. The regulatory trends noted for Assorted_Epithelial cells in different miRNAs, such as hsa-miR-126-3p, hsa-miR-222-3p, hsa-miR-223-5p, and hsa-miR-342-3p, reflect the possible involvement of structural cells in the airways in modulating the cellular reaction to the exacerbation, even though none of these signals had statistical significance through bootstrapping (p = 0.054-0.658). The inconsistent regulatory trends for these miRNAs indicate the presence of both compensatory repressive and inflammatory promotion in epithelial cells when responding to the exacerbation process. The conflicting regulatory trends for these miRNAs imply that epithelial cells experience compensatory repression (Regulation Strength > 0) and inflammatory promotion (Regulation Strength < 0), which aligns with their dual function in immune surveillance and damage sites in asthma [31].

The striking effects observed in macrophages, monocytes, and dendritic cells further demonstrate that immune cells are the main downstream effectors of blood-borne miRNAs. The positive Regulation Strength values seen in these cellular fractions, specifically in cases involving hsa-miR-223-5p and hsa-miR-342-3p in DC_Assorted and Monocytes, suggest a preference for repressing transcripts that are downregulated in asthma, implying regulation to inhibit inflammation during exacerbation. Importantly, hsa-miR-222-3p exhibited a directional difference in Regulation Strength, being positive for Macrophage_Mature but negative for Macrophage_Alveolar_CC, which implies that miRNA regulation is not consistent even within a specific cell line, but depends on the differentiation status of macrophages. More modest Regulation Strength scores in the subpopulations of lymphocytes suggest more regulated or lineage-specific effects. In contrast, B_Plasma cells did not exhibit any statistically significant regulatory influence for any of the miRNAs (all p = 1.000), suggesting that this cellular subset is largely unaffected by the regulatory influence exerted by circulating miRNAs discussed in this study.

The realization that a single miRNA might regulate in opposite ways between an epithelial population and an immune population (such as miR-223-5p or miR-342-3p) highlights the importance of performing regulatory inferences on a cell type-specific basis. Measurements of circulating miRNAs through bulk techniques cannot account for such a regulation [32]. This supports the idea that circulating miRNAs play a role in the progression of the disease by re-distributing the regulatory effect between airway structural and immune cells [33].

### 4.3 Biological implications for airway inflammation and exacerbation susceptibility

Inferred regulatory interactions correspond closely to known immunological pathways. Decreased inhibition of miR-222-3p in dendritic cells suggests increased presentation of antigens and T-cell stimulation, and reduced suppression of miR-342-3p in macrophages correlates well with elevated IL-6 and TNF production [34, 35]. The increase in repression via miR-126-3p and miR-146b-5p in epithelial cells and immune cells could indicate an anti-inflammatory response designed to maintain barrier stability or prevent excessive inflammation [36, 37].

### 4.4 Independent validation and alignment with differential expression patterns

Based on GEO datasets that are independent from our predictions, we were able to verify that certain of the identified circulating miRNAs show differential expression among the pertinent immune and airway cells. These miRNAs were originally classified based on their association with exacerbation risk in the CAMP cohort [13]. For instance, miR-126-3p and miR-146b-5p, which were inversely associated with exacerbation risk, showed increased levels in healthy subjects across the GEO datasets, consistent with their proposed roles in sustaining epithelial homeostasis and inflammatory response control. On the other hand, miR-206 and miR-30e-3p, which were positively associated with exacerbation risk, showed increased levels in asthma samples. Context-dependent changes were also noted among certain miRNAs like miR-342-3p.

### 4.5 Comparison with existing computational approaches

Various approaches for miRNA functional inference from single-cell data have been described, each with their own strengths and weaknesses. One such approach is miReact [38], which estimates miRNA activity using motif enrichment on ranked gene expression profiles and is relatively simple, annotation-independent, and broadly generalizable. However, it provides no information about the role of individual target genes or the regulation direction and can only provide scalars representing miRNA activities rather than condition-specific measurements of regulation. Another method, miRSCAPE [11], uses supervised learning models trained on bulk miRNA-mRNA profiles and achieves high precision in predicting relative miRNA abundance in cancer patient data sets. Nevertheless, it requires paired and tissue-matched bulk RNA profiles and cannot measure condition-specific regulation direction. Finally, miTEA-HiRes [39] improves the resolution of enrichment approaches but, similar to other methods of this category, assigns equal weight to all target genes in the process, failing to account for different binding affinity or regulation strength and direction.

However, none of these methods are tailored for circulating miRNAs and their impact on distant cell populations. In this work, we overcome the limitations of existing methods by combining interaction evidence for miRNAs and targets, computing probability weighting for their interactions, building patient-specific models for each cell type based on single-cell RNA-sequencing data, and using SHAP interpretation to assess directionality of miRNA effects at the level of genes. As a result, we obtain Regulation Strength score that measures not only effect size but also directionality of miRNA regulation.

### 4.6 Limitations and future Directions

There are a number of limitations with this study. First, while SHAP-based regulation estimates allow for directional inference, they do not allow us to draw any conclusions regarding causality. Moreover, the directions of miRNA effects must be experimentally validated.

In our study, we concentrated on a particular set of exacerbation-specific miRNAs that had been characterized within the CAMP study group[13]. Even though our method is applicable to any miRNA panel, we concentrated specifically on this panel because we were more interested in the biology of miRNAs that have associations with exacerbations, which are one of the major sources of morbidity among patients with asthma. Thus, there might be some other circulating miRNAs that have not been included in the CAMP panel.

Another limitation concerns our use of differentially expressed genes between asthma and control sputum samples to identify cell-type-specific target genes. The scRNA-seq dataset was not a large sample size and may have impacted our ability to detect differential expression of genes, especially in less prevalent cell populations, resulting in the inability to detect relevant miRNA targets. However, in choosing to employ the DEGs in an asthma vs control comparison rather than just looking at gene expression in asthma samples only, we wanted to be able to look at the specific miRNA targeting activity associated with the asthma disease state and not the cell-type gene expression; nevertheless, our results are inevitably linked to those DEGs that were identified in our study and a greater sample size would allow for new insights into the regulatory interactions.

In addition, clustering cells can also lead to transcriptional heterogeneity being masked. Smaller cohort sizes could make our results less generalizable. Further research using larger cohorts and longitudinal analysis will help resolve these issues.

## 5 Conclusions

In this paper, we propose an approach to studying regulatory effects of circulating miRNAs in relation to specific airway and immune cells asthma exacerbations using single cell-resolved data. Through target modeling, SHAP influence analysis, and Regulation Strength estimation, we showed that eight circulating miRNAs exhibit differential effects on specific cell types, where the strongest and most statistically significant signals were observed among innate immune cells specifically monocytes, dendritic cells, and macrophages subtypes. Other epithelial cell type was shown to have directionally significant regulatory effect for several miRNAs such as hsa-miR-126-3p, hsa-miR-222-3p, hsa-miR-223-5p, and hsa-miR-342-3p. However, this effect was insignificant when assessed by bootstrap procedure. In addition, the effect of hsa-miR-222-3p in macrophage subtypes is contradictory, indicating that the effects of the miRNA can differ depending on the subtype within the lineage, while there was no regulatory effect of any of the studied miRNAs for B_Plasma cells. The consistency of these results was validated using independent GEO datasets, demonstrating high expression of protective miRNAs such as hsa-miR-126-3p, and hsa-miR-146b-5p in healthy control subjects and high expression of exacerbation-related miRNAs like hsa-miR-206 in asthma samples.

