## Supplementary Materials for "Asthma Exacerbations: Integrative Analysis of miRNA Activity Using Single-Cell Transcriptomics"

### Supplementary Methods

#### S1. Circulating miRNA Panel

A panel of eight circulating miRNAs previously associated with asthma exacerbations was curated from a serum-based study of the Childhood Asthma Management Program (CAMP) cohort [1]. In that study, circulating miRNA expression was profiled in serum samples from children, and logistic regression analysis was used to identify miRNAs predictive of future exacerbations. Exacerbation was defined as experiencing more than one systemic corticosteroid burst during the one-year follow-up period after randomization to inhaled corticosteroid treatment. The selected miRNAs were: hsa-miR-206, hsa-miR-223-5p, hsa-miR-146b-5p, hsa-miR-342-3p, hsa-miR-222-3p, hsa-miR-30e-3p, hsa-miR-126-3p, and hsa-miR-126-5p.

#### S2. miRNA Target Identification

Experimentally validated miRNA-gene interactions were retrieved using the multiMiR R package [2], which integrates data from miRecords, TarBase, and miRTarBase. These databases compile interactions supported by experimental techniques such as luciferase reporter assays, CLIP-based methods, qRT-PCR, and Western blotting. To ensure high-confidence interactions, a gene was retained as a target only if it was supported by at least two independent databases or two distinct experimental methods.

#### S3. Single-Cell RNA-Seq Data and Preprocessing

Single-cell RNA sequencing (scRNA-seq) data were obtained from induced sputum samples of 16 asthma patients and 8 healthy controls [3]. A total of 37,565 cells were profiled, with a median of 1,203 detected genes and an average of 4,186 unique molecular identifiers (UMIs) per cell. Clustering and integration analysis were performed using Seurat [4], resulting in 28 clusters that were subsequently annotated into 19 cell populations based on canonical marker genes and correlation with the Human Primary Cell Atlas [5]. These populations included major myeloid and lymphoid cell types, as well as epithelial cells.

Differential gene expression (DEG) between asthma and control samples was computed for each cell type using the `FindMarkers` function in Seurat. Genes were tested if expressed in at least 25% of cells in either group (`min.pct = 0.25`), with a minimum $\log_2$ fold change threshold of 0.25. Statistical significance was defined as $p < 0.05$ and $\log_2 \mathrm{FC} > 0.5$.

### S4. Mapping miRNA Targets to Cell-Type-Specific DEGs

For each miRNA, validated target genes were intersected with cell-type-specific DEGs. The resulting set of shared targets represents candidate genes through which circulating miRNAs may exert regulatory effects in specific cell types.

### S5. miRNA Binding Weights and Repression Score

Binding probabilities for miRNA-gene pairs were obtained from the miRWalk 3.0 database [6] and used as gene-level regulatory weights. Missing values were assigned a weight of zero. Since mature miRNAs cannot be quantified using scRNA-seq, their activity was inferred from the expression of their target genes, such that high miRNA activity corresponds to low target gene expression, and low miRNA activity corresponds to high target gene expression. For each cell, a miRNA repression score was computed as:

$$\text{Repression Score} = -\sum(\text{gene expression} \times \text{binding weight})$$

The negative sign reflects the canonical repressive function of miRNAs. This score served as a proxy for miRNA activity at single-cell resolution and was used as the response variable in downstream modeling.

### S6. Elastic Net Modeling

To model the relationship between gene expression and miRNA regulatory activity, we employed Elastic Net regression [7], which combines $L_1$ and $L_2$ regularization to handle multicollinearity and perform feature selection. For each miRNA-cell type pair, the repression score was used as the response variable, and standardized expression levels of shared target genes were used as predictors. Models were trained using five-fold cross-validation, exploring a range of regularization parameters ($\alpha \in [0,1]$ in increments of 0.1, and a logarithmically spaced grid of $\lambda$ values) to identify the optimal configuration. Genes with nonzero coefficients in the final model were considered key contributors to miRNA-mediated regulation.

### S7. SHAP-Based Feature Contribution Analysis

To interpret model predictions, SHapley Additive exPlanations (SHAP) [8] were applied to quantify the contribution of each gene to the predicted repression score. SHAP values provide a consistent, gene-level measure of feature importance across cells.

### S8. Directional Inference of miRNA Activity

SHAP values were combined with differential expression analyses to determine whether a miRNA was more or less active in asthma, in terms of increased or decreased repression of its target genes. Genes upregulated in asthma with positive SHAP values indicate low levels of repression by the miRNA. Conversely, genes downregulated in asthma with positive SHAP values indicate high levels of repression by the miRNA.

### S9. Regulation Strength Metric

To quantify the magnitude and direction of miRNA regulation within each cell type, we defined a Regulation Strength metric. For each miRNA $m$ and cell type $c$, the metric integrates SHAP-derived gene contributions with differential expression direction:

$$
\begin{aligned}
S_{\text{up}} &= \sum_{g \in G_c, \text{DE}_g = +1} |\text{SHAP}_g|, \\
S_{\text{down}} &= \sum_{g \in G_c, \text{DE}_g = -1} |\text{SHAP}_g|, \\
R_{m,c} &= S_{\text{down}} - S_{\text{up}}
\end{aligned}
$$

where $G_c$ is the set of shared target genes with available SHAP values in cell type $c$, and $\text{DE}_g \in \{+1, -1\}$ denotes the direction of differential expression for gene $g$ in asthma. A positive $R_{m,c}$ indicates preferential repression of asthma-downregulated genes, suggesting increased miRNA activity, while a negative $R_{m,c}$ indicates preferential repression of asthma-upregulated genes, suggesting reduced miRNA activity. The magnitude of $R_{m,c}$ reflects the strength of this directional bias.

### S10. Bootstrap Significance Testing

To assess the statistical robustness of each Regulation Strength estimate, we performed cell-level bootstrap resampling. For each miRNA-cell-type pair, cells were resampled with replacement 1,000 times, and $R_{m,c}$ was recomputed for each resampled iteration to generate an empirical distribution. From this distribution, 95% confidence intervals were derived, and a two-sided bootstrap $p$-value was computed as the proportion of resampled iterations for which the sign of $R_{m,c}$ disagreed with the observed sign (or, equivalently, the proportion of the resampled distribution crossing zero, doubled). Pairs with bootstrap $p < 0.05$ were considered statistically significant. Complete results for all miRNA-cell-type pairs are provided in Supplementary Table 1.

### S11. Permutation-Based Single-Cell Significance Maps

To evaluate the statistical robustness of miRNA activity at the level of individual cells, a null distribution was generated for each cell by permuting the association between target genes and their SHAP-derived weights while holding gene expression fixed. Observed miRNA activity was compared against this null distribution, and resulting $p$-values were transformed to $-\log_{10}(p)$ for visualization, with higher values indicating stronger evidence of non-random miRNA regulation (see Fig. 2C-D in the main text).

### S12. External Validation Using Independent GEO Datasets

To validate the inferred regulatory relationships, four independent GEO datasets were analyzed: GSE120172 [9] (pediatric dust-mite asthma; bronchial epithelial cells and plasma), GSE61563 [10] (chronic asthma; bronchial epithelial cells), GSE142237 [11, 12] (steroid-naive asthma; bronchial epithelial cells), and GSE146306 [13] (severe asthma; sputum and immune cells). Each dataset was downloaded and processed using the GEOquery package [14], and expression matrices were filtered to retain the eight candidate miRNAs. Samples were classified into asthma or healthy control groups using accompanying phenotype metadata, and expression values were log-transformed where applicable.

Differential expression between asthma and control groups was evaluated using the limma framework [15]. A design matrix representing group status was constructed, linear models were fitted using `lmFit`, and variance was stabilized using empirical Bayes moderation (`eBayes`). For each miRNA, $\log_2$ fold changes and Benjamini-Hochberg-adjusted $p$-values [16] were computed. Consistent direction and magnitude of differential expression across independent datasets were interpreted as supporting evidence for the predicted miRNA-cell-type regulatory relationships.

### S13. Software and Package Versions

All analyses were performed in R and Python. Key packages included: Seurat (single-cell clustering and DEG analysis), multiMiR (miRNA-target retrieval), miRWalk 3.0 (binding probability weights), scikit-learn (Elastic Net regression, five-fold cross-validation), SHAP (Python package, gene-level feature attribution), GEOquery and limma (GEO dataset processing and differential expression), and UMAP (dimensionality reduction and visualization).

**Supplementary Table 1.** Bootstrap confidence intervals (95%, n=1,000 iterations) for Regulation Strength across all miRNA–cell type pairs.

| miRNA | Cell Type | Observed | 95% CI | p |
|---|---|---|---|---|
| hsa-miR-206 | DC_Assorted | +0.1866 | [+0.1157, +0.2568] | 0.0010* |
| hsa-miR-222-3p | DC_Assorted | +0.3486 | [+0.2321, +0.4708] | 0.0010* |
| hsa-miR-223-5p | DC_Assorted | +0.1689 | [+0.1013, +0.2390] | 0.0010* |
| hsa-miR-342-3p | DC_Assorted | +0.1904 | [+0.1341, +0.3000] | 0.0010* |
| hsa-miR-222-3p | Macrophage_Mature | +0.2449 | [+0.1936, +0.3174] | 0.0010* |
| hsa-miR-126-3p | Monocyte | +0.3104 | [+0.2593, +0.3617] | 0.0010* |
| hsa-miR-146b-5p | Monocyte | +0.3502 | [+0.2874, +0.4113] | 0.0010* |
| hsa-miR-206 | Monocyte | +0.2598 | [+0.2229, +0.3052] | 0.0010* |
| hsa-miR-223-5p | Monocyte | +0.1132 | [+0.0871, +0.1567] | 0.0010* |
| hsa-miR-30e-3p | Monocyte | +0.2323 | [+0.1894, +0.2724] | 0.0010* |
| hsa-miR-126-5p | T | −0.1076 | [−0.1601, −0.0515] | 0.0010* |
| hsa-miR-206 | T | +0.1000 | [+0.0660, +0.1743] | 0.0010* |
| hsa-miR-223-5p | T | +0.1325 | [+0.0689, +0.1906] | 0.0010* |
| hsa-miR-30e-3p | T | +0.1007 | [+0.0390, +0.1611] | 0.0010* |
| hsa-miR-342-3p | T | +0.3164 | [+0.2187, +0.4224] | 0.0010* |
| hsa-miR-126-5p | T_Cytotoxic | +0.3444 | [+0.2580, +0.4257] | 0.0010* |
| hsa-miR-206 | T_Cytotoxic | +0.2229 | [+0.0861, +0.3552] | 0.0010* |
| hsa-miR-223-5p | T_Cytotoxic | +0.2029 | [+0.0908, +0.3622] | 0.0020* |
| hsa-miR-146b-5p | T_Cytotoxic | +0.0526 | [+0.0200, +0.0897] | 0.0030* |
| hsa-miR-146b-5p | DC_Assorted | +0.1330 | [+0.0604, +0.2383] | 0.0040* |
| hsa-miR-342-3p | Monocyte | +0.0764 | [+0.0320, +0.1354] | 0.0040* |
| hsa-miR-222-3p | Macrophage_Alveolar_CC | −0.3078 | [−0.6345, −0.1747] | 0.0150* |
| hsa-miR-342-3p | Assorted_Epithelial | −0.0402 | [−0.0979, −0.0162] | 0.0540 |
| hsa-miR-342-3p | B | +0.0746 | [+0.0300, +0.1897] | 0.0610 |
| hsa-miR-223-5p | Macrophage_Alveolar_CC | −0.2260 | [−0.5643, −0.0675] | 0.0660 |
| hsa-miR-126-3p | Assorted_Epithelial | −0.0319 | [−0.0892, −0.0079] | 0.1260 |
| hsa-miR-126-3p | B | +0.0108 | [+0.0006, +0.0356] | 0.2210 |
| hsa-miR-223-5p | Assorted_Epithelial | −0.0316 | [−0.1125, −0.0111] | 0.2240 |
| hsa-miR-126-3p | T_Cytotoxic | +0.0450 | [+0.0021, +0.1380] | 0.2270 |
| hsa-miR-126-5p | ILC_Assorted | −0.0754 | [−0.2673, −0.0037] | 0.3330 |
| hsa-miR-30e-3p | Macrophage_Alveolar_CC | −0.0359 | [−0.1942, −0.0032] | 0.5160 |
| hsa-miR-126-3p | ILC_Assorted | +0.0139 | [+0.0082, +0.1377] | 0.6790 |
| hsa-miR-223-5p | Macrophage_Mature | +0.0024 | [+0.0005, +0.0330] | 0.8200 |
| hsa-miR-30e-3p | B | −0.0024 | [−0.1727, −0.0018] | 0.9580 |

*Continued on next page*

Supplementary Table 1 – *continued from previous page*

| miRNA | Cell Type | Observed | 95% CI | p |
|---|---|---|---|---|
| hsa-miR-206 | ILC_Assorted | +0.0004 | [+0.0012, +0.0889] | 0.9830 |
| hsa-miR-146b-5p | B_Plasma | −0.0000 | [−0.1618, −0.0034] | 1.0000 |
| hsa-miR-206 | B_Plasma | −0.0000 | [−0.2232, −0.0119] | 1.0000 |
| hsa-miR-223-5p | B_Plasma | −0.0000 | [−0.2874, −0.0135] | 1.0000 |
| hsa-miR-30e-3p | B_Plasma | −0.0000 | [−0.1089, −0.0008] | 1.0000 |
| hsa-miR-222-3p | Assorted_Epithelial | +0.2154 | [−0.0336, +0.4725] | 0.1000 |
| hsa-miR-222-3p | T_Cytotoxic | +0.1696 | [−0.0319, +0.3986] | 0.1160 |
| hsa-miR-206 | Macrophage_Mature | −0.0082 | [−0.0226, +0.0074] | 0.3070 |
| hsa-miR-222-3p | Monocyte | +0.0182 | [−0.0136, +0.0684] | 0.3380 |
| hsa-miR-206 | B | +0.0251 | [−0.0177, +0.0964] | 0.3970 |
| hsa-miR-222-3p | Macrophage_Immature | −0.0083 | [−0.0255, +0.0164] | 0.4330 |
| hsa-miR-146b-5p | T | −0.0299 | [−0.0954, +0.0484] | 0.4330 |
| hsa-miR-222-3p | T_Regulatory | +0.0673 | [−0.0011, +0.4990] | 0.6070 |
| hsa-miR-342-3p | ILC_Assorted | +0.0592 | [−0.1131, +0.3294] | 0.6100 |
| hsa-miR-206 | Assorted_Epithelial | −0.0151 | [−0.0770, +0.0462] | 0.6320 |
| hsa-miR-30e-3p | Assorted_Epithelial | +0.0481 | [−0.1019, +0.2791] | 0.6580 |
| hsa-miR-222-3p | T | +0.0149 | [−0.1301, +0.1572] | 0.8540 |
| hsa-miR-222-3p | ILC_Assorted | −0.0162 | [−0.3382, +0.5654] | 0.9280 |
| hsa-miR-342-3p | Macrophage_Alveolar_CC | −0.0042 | [−0.1456, +0.1032] | 0.9340 |
| hsa-miR-342-3p | T_Cytotoxic | +0.0036 | [−0.1166, +0.1386] | 0.9650 |
| hsa-miR-223-5p | ILC_Assorted | −0.0027 | [−0.0947, +0.1245] | 0.9670 |
| hsa-miR-222-3p | B | −0.0021 | [−0.1949, +0.1975] | 0.9850 |
| hsa-miR-30e-3p | T_Cytotoxic | +0.0002 | [−0.0773, +0.0756] | 0.9960 |
| hsa-miR-223-5p | B | −0.0000 | [−0.0000, +0.0000] | 1.0000 |
| hsa-miR-126-5p | B_Plasma | −0.0000 | [−0.0000, +0.0000] | 1.0000 |
| hsa-miR-222-3p | B_Plasma | −0.0000 | [−0.6834, +0.3520] | 1.0000 |
| hsa-miR-342-3p | B_Plasma | +0.0000 | [−0.0169, +0.4289] | 1.0000 |
| hsa-miR-126-5p | Macrophage_Alveolar_CC | +0.0000 | [+0.0000, +0.0000] | 1.0000 |
| hsa-miR-342-3p | Macrophage_Immature | +0.0000 | [+0.0000, +0.0000] | 1.0000 |
| hsa-miR-222-3p | Mast | +0.0000 | [−0.0620, +0.6857] | 1.0000 |
| hsa-miR-223-5p | Mast | +0.0000 | [+0.0000, +0.0000] | 1.0000 |
| hsa-miR-126-5p | Monocyte | +0.0000 | [+0.0000, +0.0000] | 1.0000 |
| hsa-miR-223-5p | Multiplet_MyeloidLymphoid | −0.0000 | [−0.0000, +0.0000] | 1.0000 |
| hsa-miR-223-5p | pDC | +0.0000 | [+0.0000, +0.0000] | 1.0000 |
| hsa-miR-342-3p | pDC | −0.0000 | [−0.3030, +0.0549] | 1.0000 |

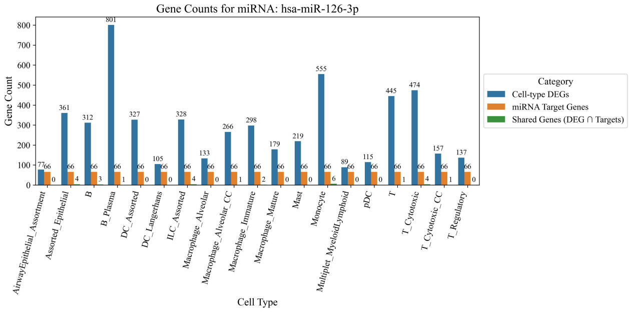


**(a)** Gene counts for hsa-miR-126-3p across cell types.

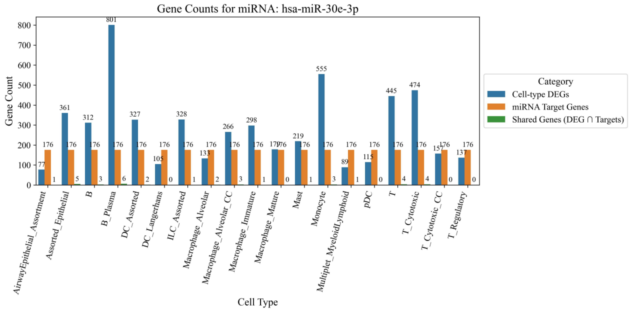


**(b)** Gene counts for hsa-miR-30e-3p across cell types.

**Figure 1.** Gene counts for differentially expressed genes (DEGs), predicted miRNA target genes, and their overlap across annotated airway and immune cell types.

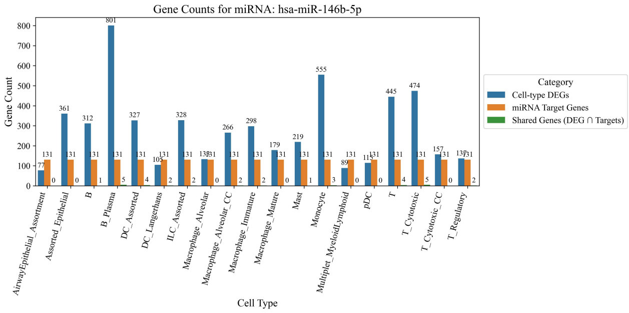


**(a)** Gene counts for hsa-miR-146b-5p across cell types.

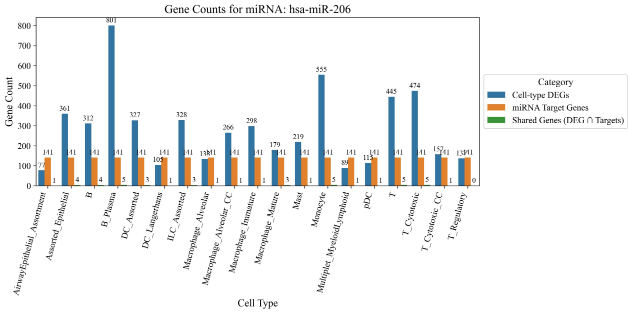


**(b)** Gene counts for hsa-miR-206 across cell types.

**Figure 2.** Gene counts for differentially expressed genes (DEGs), predicted miRNA target genes, and their overlap across annotated airway and immune cell types.

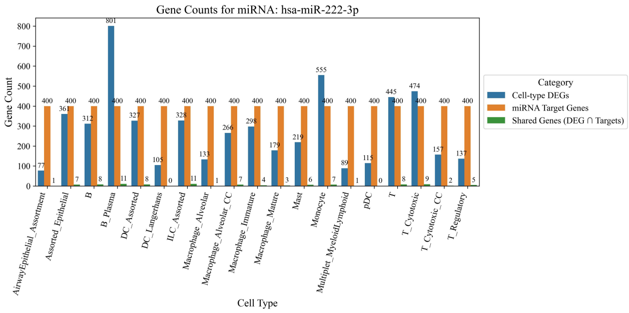


**(a)** Gene counts for hsa-miR-222-3p across cell types.

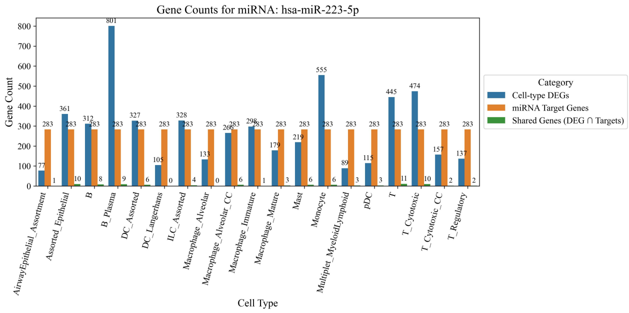


**(b)** Gene counts for hsa-miR-223-5p across cell types.

**Figure 3.** Gene counts for differentially expressed genes (DEGs), predicted miRNA target genes, and their overlap across annotated airway and immune cell types.

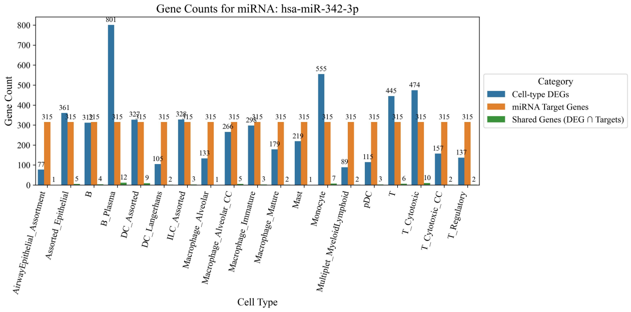


**(a)** Gene counts for hsa-miR-342-3p across cell types.

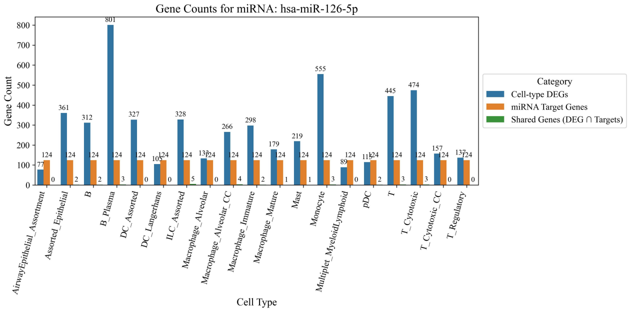


**(b)** Gene counts for hsa-miR-126-5p across cell types.

**Figure 4.** Gene counts for differentially expressed genes (DEGs), predicted miRNA target genes, and their overlap across annotated airway and immune cell types.

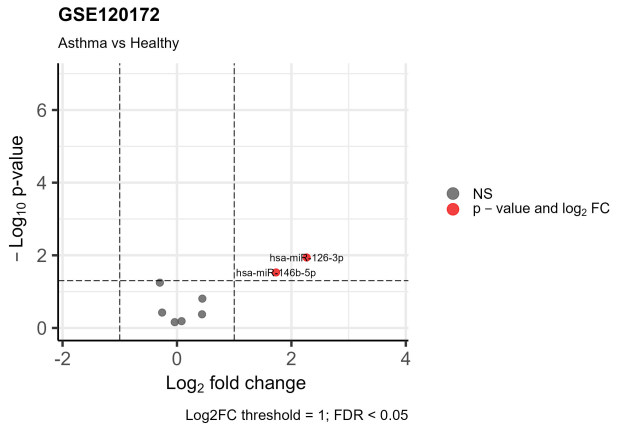


**(a)** Volcano plot of differentially expressed miRNAs in GSE120172 (Asthma vs. Healthy).

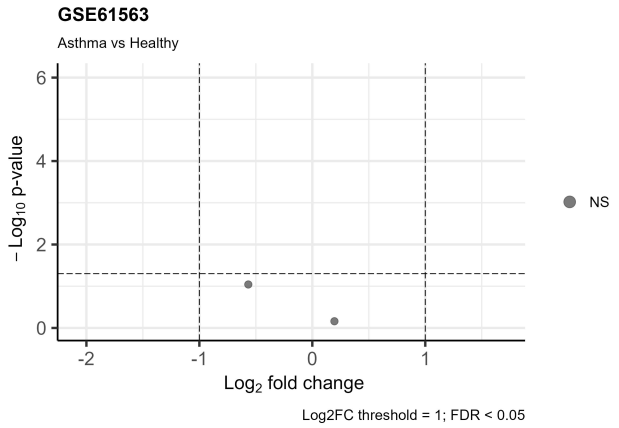


**(b)** Volcano plot of differentially expressed miRNAs in GSE61563 (Asthma vs. Healthy).

**Figure 5.** Volcano plots showing differential miRNA expression for two GEO datasets.

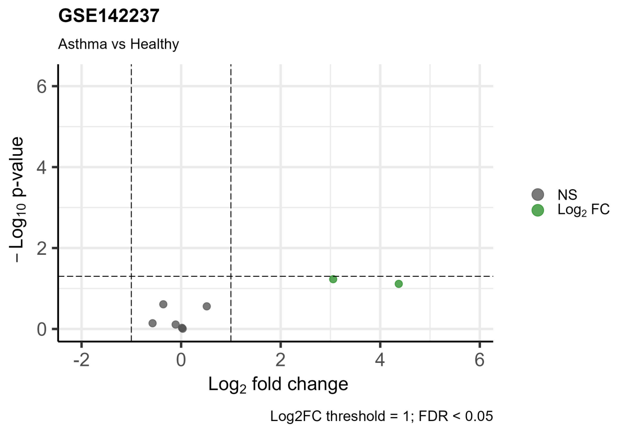


**(a)** Volcano plot of differentially expressed miRNAs in GSE142237 (Asthma vs. Healthy).

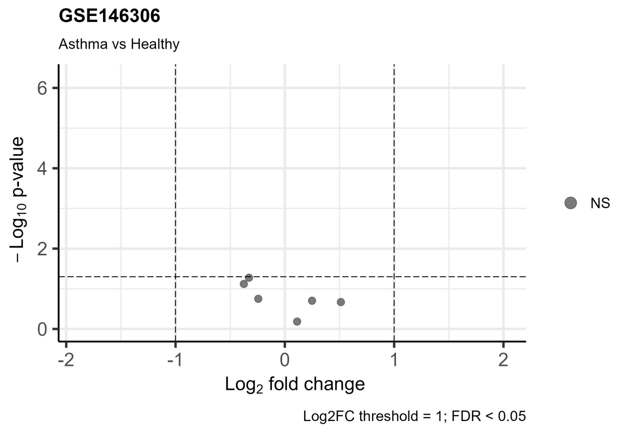


**(b)** Volcano plot of differentially expressed miRNAs in GSE146306 (Asthma vs. Healthy).

**Figure 6.** Volcano plots showing differential miRNA expression for two GEO datasets.

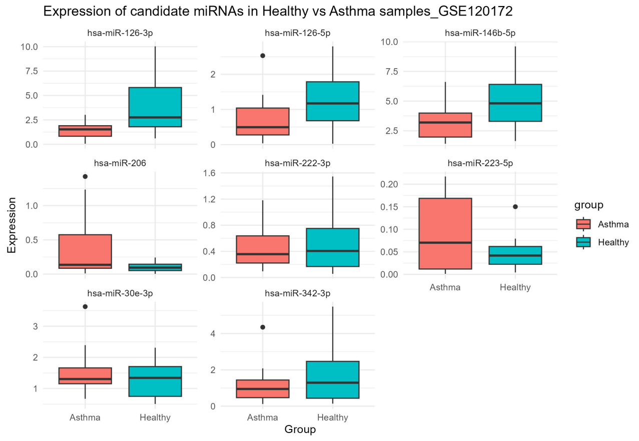


**(a)** Expression of candidate circulating miRNAs in asthma and healthy samples from GSE120172.

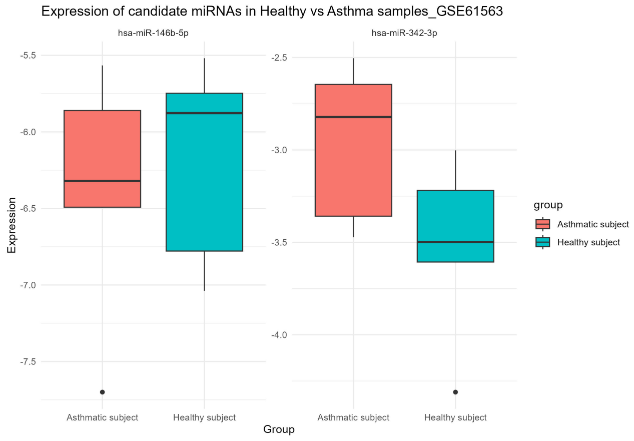


**(b)** Expression of candidate circulating miRNAs in asthma and healthy samples from GSE61563.

**Figure 7.** Expression of candidate circulating miRNAs in asthma versus healthy samples for two GEO datasets.

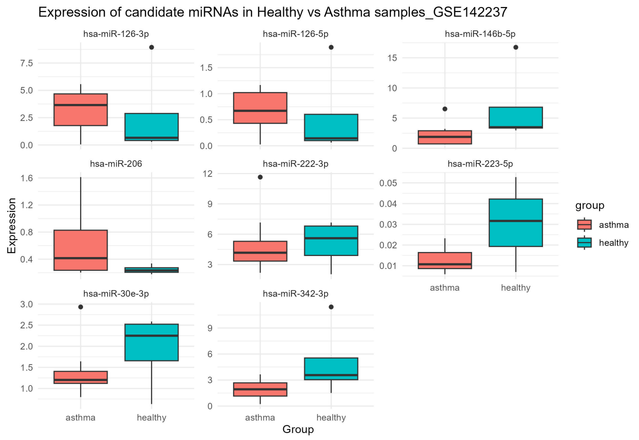


**(a)** Expression of candidate circulating miRNAs in asthma and healthy samples from GSE142237.

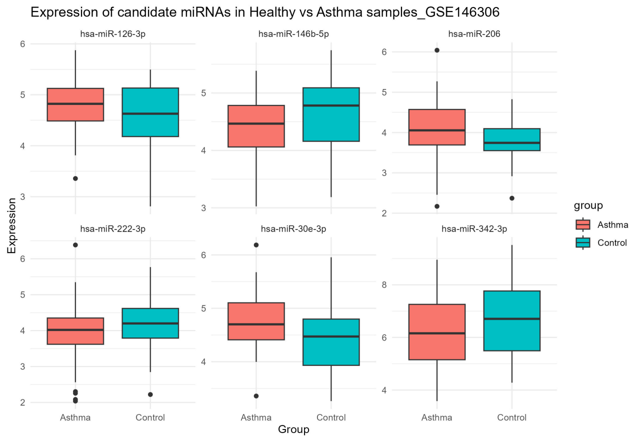


**(b)** Expression of candidate circulating miRNAs in asthma and healthy samples from GSE146306.

**Figure 8.** Expression of candidate circulating miRNAs in asthma versus healthy samples for two GEO datasets.

### Funding

This work was supported by the National Institutes of Health (NIH) grants R01 HL162570, R01 HL161362, R01 HL155742, R01 HL177625 and K99 HL183694.
